# A colorectal cancer organoid-CAF co-culture chip for drug screening and personalized therapy

**DOI:** 10.64898/2026.06.15.732195

**Authors:** Rongrong Xiao, Zhijie Wang, Yu Zhou, Duanchen Ding, Jian Wang, Jianchuang Liu, Yi Wang, Qian Liu, Xiaoni Ai

## Abstract

The tumor microenvironment (TME) plays a critical role in cancer progression and therapeutic response, with cancer-associated fibroblasts (CAFs) being a key stromal component. Conventional tumor organoid models lack TME elements, and existing co-culture systems have limitations in recapitulating dynamic, multidimensional interactions. Here, we developed a novel dynamic co-culture chip (BAC) to better model the TME and investigate CAF-tumor cell crosstalk. Using this platform, we established a non-contact co-culture model of patient-derived colorectal cancer cells and CAFs. The resulting model closely recapitulated the morphological and molecular characteristics of the original patient tumors. Compared with tumor organoids cultured alone, the co-culture system exhibited significantly higher resistance to the clinically common chemotherapeutics 5-fluorouracil and oxaliplatin. Moreover, the presence of CAFs promoted tumor recurrence. Notably, drug responses in the co-culture model showed superior concordance with clinical outcomes relative to both organoid-only and animal models. Transcriptomic profiling under different culture conditions provided further insights into the mechanisms driving CAF-mediated interactions. These findings demonstrate that the BAC-based tumor organoid-CAF co-culture model serves as a more accurate platform for predicting drug responses, investigating TME-dependent mechanisms, and guiding personalized cancer therapy.

## 1. Introduction

The complexity of the tumor microenvironment (TME) poses a major challenge in cancer therapy. Within this microenvironment, in addition to tumor cells themselves, various other cell type, such as fibroblasts, immune cells, and endothelial cell, interact intricately to influence tumor growth, invasion, and treatment response.

For a long time, various disease research models have served as essential tools in biomedical research. Two- or three-dimensional cell line models and animal models have been the primary systems for modeling human development and disease ^[1]^. With the emergence and development of tumor organoid technology, it has become an important in vitro model for anticancer drug development and personalized medicine. Tumor organoids are mainly derived from surgically resected tumor specimens, biopsy tissues, or liquid biopsy samples. After appropriate processing, they form three-dimensional cell clusters with proliferative and self-organizing capacity under specific culture conditions in vitro ^[2]^. These organoids can be expanded long-term, cryopreserved, and genetically modified, while maintaining genetic and phenotypic stability. Based on these advantages, tumor organoids have been widely used in studies of tumor pathogenesis ^[3]^, drug screening ^[4]^, target discovery ^[5]^, personalized medicine ^[6]^, and other related fields.

However, current tumor organoid models often lack this crucial microenvironment, limiting their predictive utility ^[7]^. As the most abundant stromal cell type in the TME, cancer-associated fibroblasts (CAFs) exert profound tumor-promoting effects through multiple mechanisms ^[8]^. Studies have shown that multiple growth factors secreted by CAFs, such as EGF, IGF, and SDF-1, can activate downstream signaling pathways and promote cancer cell proliferation ^[9]^. In addition, CAFs play a crucial role in regulating the immune response against tumors. CAFs can promote the expression of immunosuppressive molecules PD-1 and CTLA-4 on activated T cells, creating an immunosuppressive microenvironment. CAFs also secrete tumor-promoting factors such as TGF-β and IL-10, which directly suppress cytotoxic T cells while inducing Tregs and MDSCs, indirectly inhibiting T cell activation ^[10]^. CAFs are also involved in drug resistance, forming a physical protective niche by secreting collagen, activating Met/PI3K/AKT signaling, and upregulating GRP78 ^[11]^. Therefore, compared with tumor organoids alone, co-culture models combining tumor organoids with CAFs have greater value for investigating tumor mechanisms and evaluating antitumor drugs.

To better understand the interactions between CAFs and tumor cells, researchers have developed various in vitro models, including traditional co-culture systems and microfluidic-based three-dimensional culture systems. Currently, scientists have established culture systems that model the TME using different approaches. *Schuth et al.* found that introducing CAFs into organoid co-cultures more faithfully recapitulates the TME; the presence of CAFs reduced the sensitivity of organoids to chemotherapeutic drugs. Using single-cell sequencing, they discovered that CAFs promote inflammation and EMT, explaining drug resistance ^[12]^. In a co-culture system of CAFs and pancreatic cancer organoids, it was shown that CAFs can provide Wnt factors for tumor cell growth ^[13]^. Pancreatic CAFs not only support tumor organoid growth but also exhibit heterogeneity, with different subpopulations playing distinct roles during tumor progression ^[14]^. Similarly, in a 3D co-culture model of CAFs and liver cancer organoids, the results demonstrated the important role of CAFs in liver cancer development and therapy resistance ^[15]^. The development of co-culture systems has also laid a foundation for studying tumors and immunotherapy. Tsai et al. were the first to establish a triple-culture model comprising tumor organoids, stromal components, and immune cells, observing CAF activation and T cell infiltration ^[16]^. However, most reported co-culture models to date are based on Transwell 2D systems or gel-droplet mixed culture systems, which rely on complex image processing and do not allow separate, multidimensional investigation of dynamic changes in cancer cells, CAFs, and the surrounding matrix, nor can they precisely decouple contact-dependent from paracrine signaling between these components.

Therefore, we have designed a novel dynamic co-culture biomimetic array chips system (BAC) that better mimics the in vivo TME and provides a powerful tool for studying interactions between CAFs and tumor cells. Based on self-designed chips, we constructed a dynamic, real-time observable, and component-separable co-culture model of tumor organoids and CAFs. Tumor tissues from colorectal cancer patients were collected, and tumor cells and fibroblasts were isolated for non-contact co-culture. The morphology and molecular characteristics of tumor cells in the model are highly consistent with those of patient tissues, and the cells exhibit higher resistance to 5-FU and oxaliplatin. Moreover, the presence of CAFs promotes tumor regrowth post-treatment. Notably, compared with tumor organoid-alone models and animal models, the drug response in the co-culture model shows higher concordance with clinical treatment outcomes. Based on these observations, we investigated gene expression profiles under different culture conditions to gain a deeper understanding of the mechanisms driving these interactions. As a more accurate drug response prediction model, we anticipate that it will better simulate the TME, more accurately reveal the cellular and molecular characteristics of tumors, thereby improving the exploration of drug-induced tumor inhibition, aiding the development of novel anticancer strategies, and providing more precise clues for patient treatment. This study aims to establish a highly predictive, modular in vitro model with physically separable components that recapitulates the TME, thereby offering a robust platform for investigating tumor–stroma crosstalk, guiding anticancer drug development, and advancing personalized cancer therapy.

## 2. Results

### 2.1 Design of dynamic co-culture chips

We have developed a set of dynamic co-culture biomimetic array chips (BAC) that enable non-contact co-culture of cancer-associated fibroblasts (CAFs) with tumor organoids, allowing investigation of cell interactions and guiding personalized medication. The BAC-OC is an organoid culture chip designed for cell expansion and sample collection after co-culture. The BAC-OD is an organoid detection chip used for drug testing on organoids and CAFs separately after co-culture (Fig. 1A and Fig. S1). Both chips are made of biocompatible polycarbonate (PC) and feature ultra-thin glass bottoms. They have standardized outer dimensions and inter-hole spacing, making them compatible with conventional pipetting and detection equipment. Top-view structural diagrams, longitudinal cross-sections, and dimensional drawings of the entire BAC chip and a single co-culture unit are shown in Fig. S2 and Fig. 1B, respectively. The BAC-OC contains 24 co-culture units. Each unit includes a central organoid culture well for 3D PDO culture and a peripheral CAF culture well for 3D CAF culture. The BAC-OD contains 48 co-culture units, each consisting of two individual culture wells (organoid well and CAF well) and a connecting region. Every cell culture well has a cell seeding area and a medium reservoir (Fig. 1B).

**Figure 1:**
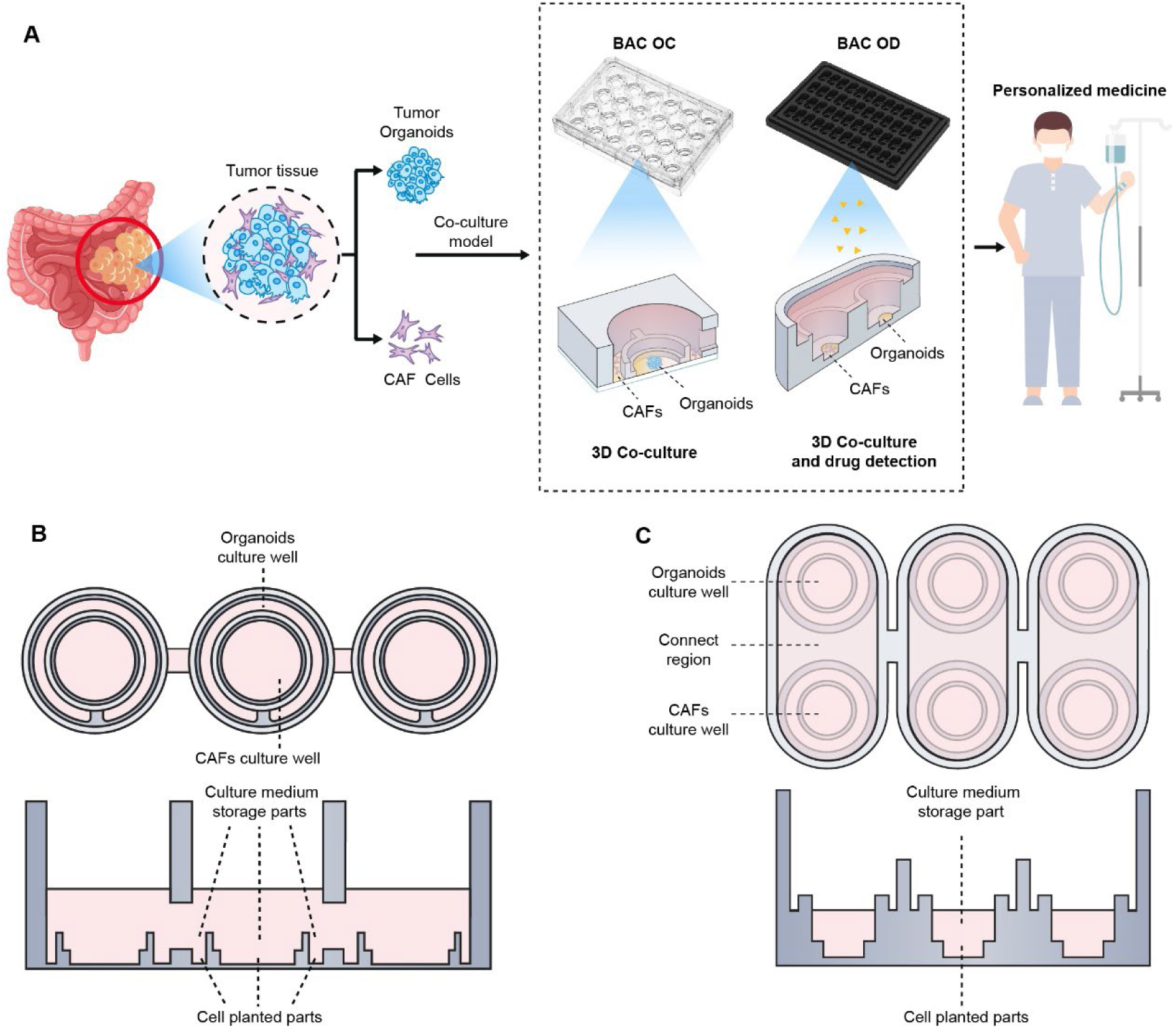
Design and Usage Details of Dynamic Co-culture Chips. A. Workflow summarizing the construction and detection of PDOs and CAFs non-contact co-culture models *in vitro* on BAC platforms. B. Schematic design details of two BAC platforms (Top view and longitudinal view).

On BAC chips, hydrogel-organoid and hydrogel-CAF mixtures are seeded into the respective loading ports. A small volume of culture medium or detection reagent is added to maintain individual cultivation of organoids and CAFs. Subsequently, co-culture is initiated by adding more medium to connect the organoid well and CAF well within a co-culture unit. If needed, the medium volume can be further increased to link three co-culture units for studying more complex multicellular or multi-organ interactions. Typically, during dynamic co-culture on the BAC-OC chip, three co-culture units are connected. Each unit is established and analyzed independently on the BAC-OD chip (Fig. 1B and Fig. S1).

### 2.2 Dynamic culture promotes growth of PDOs on chips

To promote rapid expansion of PDOs in each culture unit on the BAC-OC, we connected the internal channels and maintained liquid flow among three co-culture units using a precision shaker to create a dynamic culture environment (Fig. 2A). Quantitative fluorescence diffusion results demonstrate differences in mass transfer efficiency under static versus dynamic conditions. Under static conditions, FITC-dextran encapsulated in hydrogel within the middle well was gradually released into the medium over the first 90 min and diffused slowly; it remained unevenly distributed across the three wells even after 3 hours. Under dynamic conditions, FITC-dextran was released and rapidly diffused to both sides due to fluid flow, and its concentration remained consistent across the three wells throughout the observation period (Fig. 2B).

**Figure 2:**
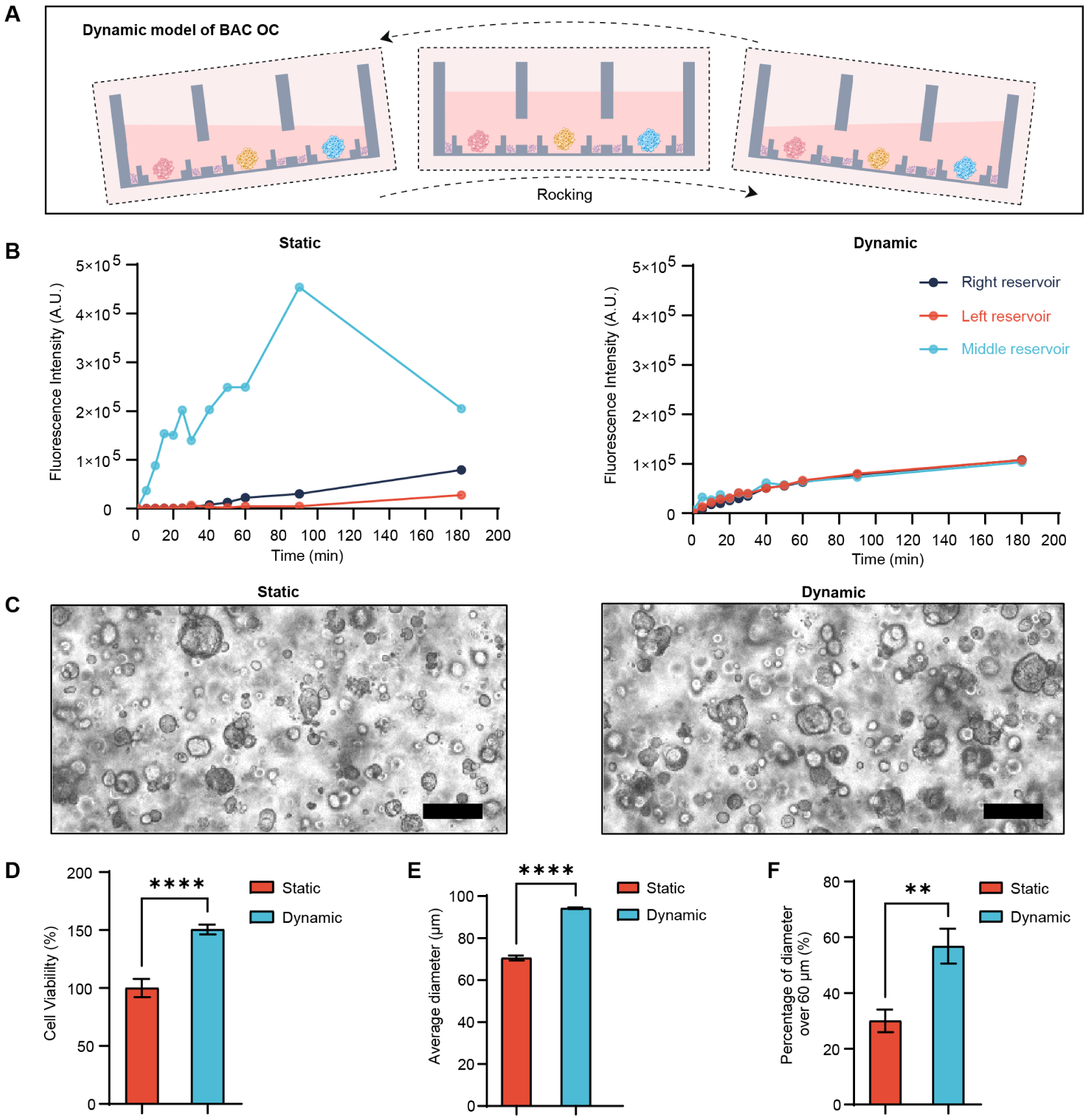
Dynamic Culture Significantly Accelerates the Growth of PDOs. A. Dynamic culture of PDOs using BAC OC by rocking. B. Characterization of molecular diffusion efficiency in dynamic and static culture models by quantifying the transfer of FITC-dextran in tandem wells of BAC OC. C. Bright field photos of dynamic PDOs and static culture PDOs. Scale bars are 100μm as indicated. D. The statistical histogram of cell viability in dynamic and static culture models. All data are presented as means ± SD of three replicates. n=6 well E. The statistical histogram of the average diameter of PDOs. F. The percentage of PDOs with a diameter over 60 μm. (**P < 0.01, ***P < 0.001; ****P < 0.0001.)

The density and size of PDOs in bright-field images were significantly larger under dynamic culture (Fig. 2C). Quantitative analysis showed that cell viability in the dynamic culture chip was significantly higher than that in static culture after the same duration (Fig. 2D). The diameter of PDOs under dynamic conditions reached 94.21 ± 0.41 μm, whereas under static conditions it was only 70.50 ± 1.18 μm (Fig. 2E). The proportion of PDOs with a diameter >60 μm increased by 26.8% compared with static culture (Fig. 2F).

Actual photos of the two chips and dynamic culture on precision shakers are shown in Fig. S3. Lateral shaking promotes nutrient exchange among three co-culture units on the BAC-OC. Vertical shaking promotes diffusion of factors between the organoid well and CAF well within a single co-culture unit while maintaining independence from the other two units (Fig. S4A). Quantitative fluorescence diffusion results on the BAC-OD demonstrate mass transfer differences under static versus dynamic conditions. Under static conditions, FITC-dextran encapsulated in the lower well diffused slowly, and only a weak fluorescent signal was detected in the upper well after 12 h. Under dynamic conditions with fluid flow, FITC-dextran concentration reached equilibrium between the lower and upper wells after about 4 h (Fig. S4B).

### 2.3 Dynamically cultured PDOs closely resemble the corresponding patient tissues

HE staining revealed heterogeneity in nuclear and cellular morphology, similar to that of epithelial components in the parental tissues, consistent with tumor cell characteristics (Fig. 3A and S5A). We confirmed the consistent expression of histology-specific markers between PDOs and parental tissues. Immunohistochemistry and immunofluorescence staining showed that PDOs retained expression of CK20, CDX2, and Ki-67 (Fig. 3A-B and S5A-B).

**Figure 3:**
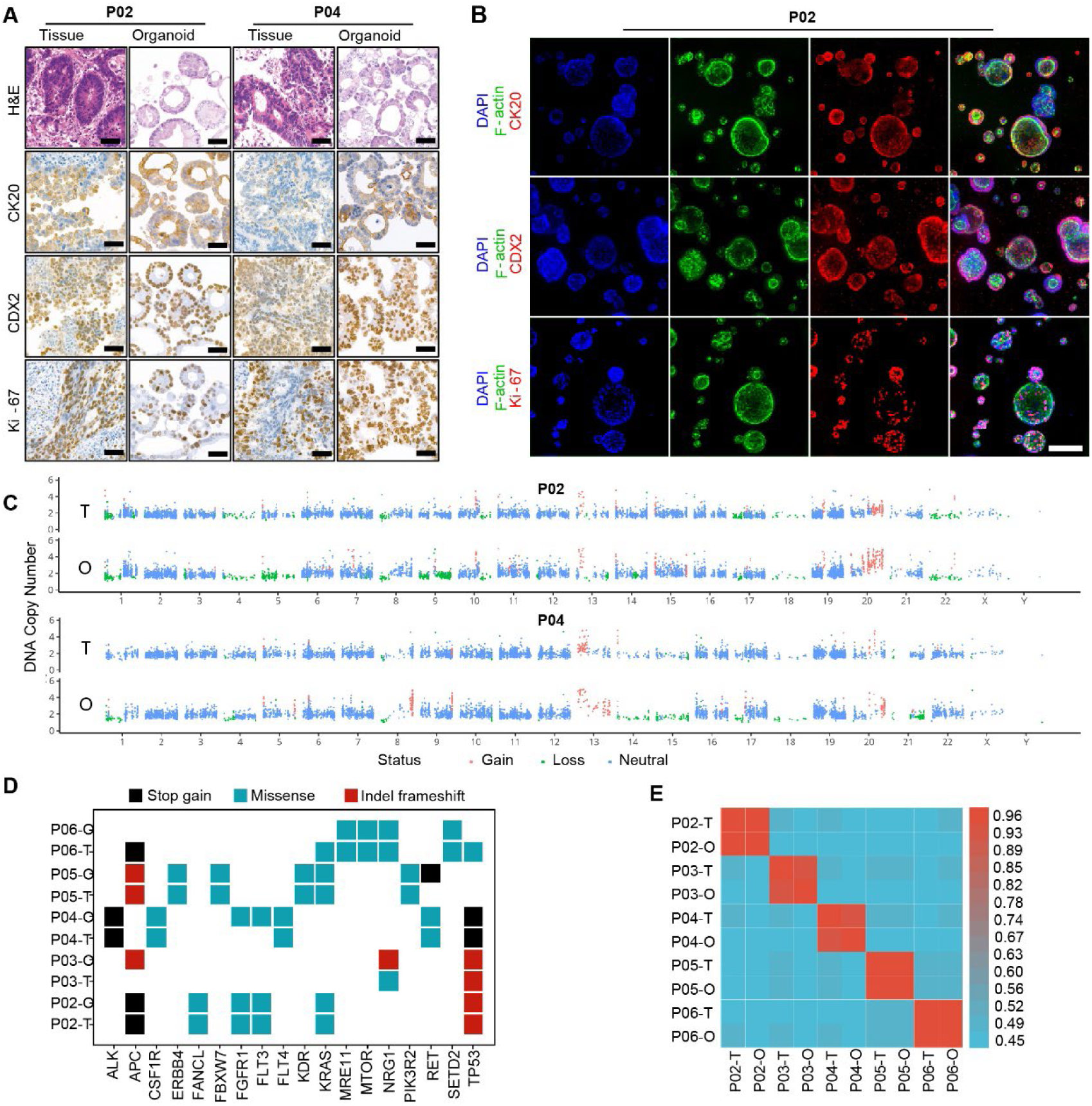
PDOs are Highly Consistent with Patient Tissues. A. Comparative H&E staining and immunohistochemical staining images of PDOs and parental tumor tissues. The typical markers are CK20, CDX2, and Ki-67. Scale bars are 50μm as indicated. B. The images of immunofluorescence staining of PDOs. The typical markers are CK20, CDX2, and Ki-67. Scale bars are 100μm as indicated. C. Scatterplots illustrating genome-wide gene CNVs of the PDOs and parental tumor tissues (red: gain, green: loss, blue: neutral). D. Heatmap showing a cancer-related set of variants identified by WES. E. Correlation heatmap between PDOs and paired tumor tissue.

To determine whether PDOs recapitulate the genomic features of original tumors, we performed whole-exome sequencing (WES) on 5 paired PDOs, tumor tissues, and matched normal tissues. The mutation profiles of PDOs were highly consistent with their parental tumors. Copy number variations (CNVs) also showed high concordance between PDOs and tumor samples (Fig. 3C and S3C). In addition, somatic mutations in driver genes such as TP53 and KRAS were observed in both tumor tissues and PDOs (Fig. 3D). Using single nucleotide variations, we generated a Pearson correlation heatmap; the correlation coefficients between PDOs and their paired tissues were all above 0.9, demonstrating that PDOs maintain the genomic features of the original tissues (Fig. 3E).

### 2.4 CAFs promote PDO growth and epithelial-mesenchymal transition (EMT) in a non-contact co-culture model

Primary CAFs were isolated from surgical tissues of colorectal cancer patients and characterized by immunofluorescence staining for specific markers. The isolated CAFs exhibited a long spindle-shaped morphology and expressed FAP, α-SMA, and VIM (Fig. 4A). PDOs were then non-contact co-cultured with paired CAFs on the BAC-OC chip. Bright-field photographs showed that CAFs significantly promoted PDO growth compared with mono-culture (Fig. 4B). The effect varied among donors; the most pronounced increase in cell viability (80.7%) was observed in donor P04 (Fig. 4C). The EMT marker fibronectin was significantly upregulated in the co-culture model compared with mono-culture (Fig. 4D and S6). Thus, non-contact co-culture with CAFs promotes PDO proliferation and EMT.

**Figure 4:**
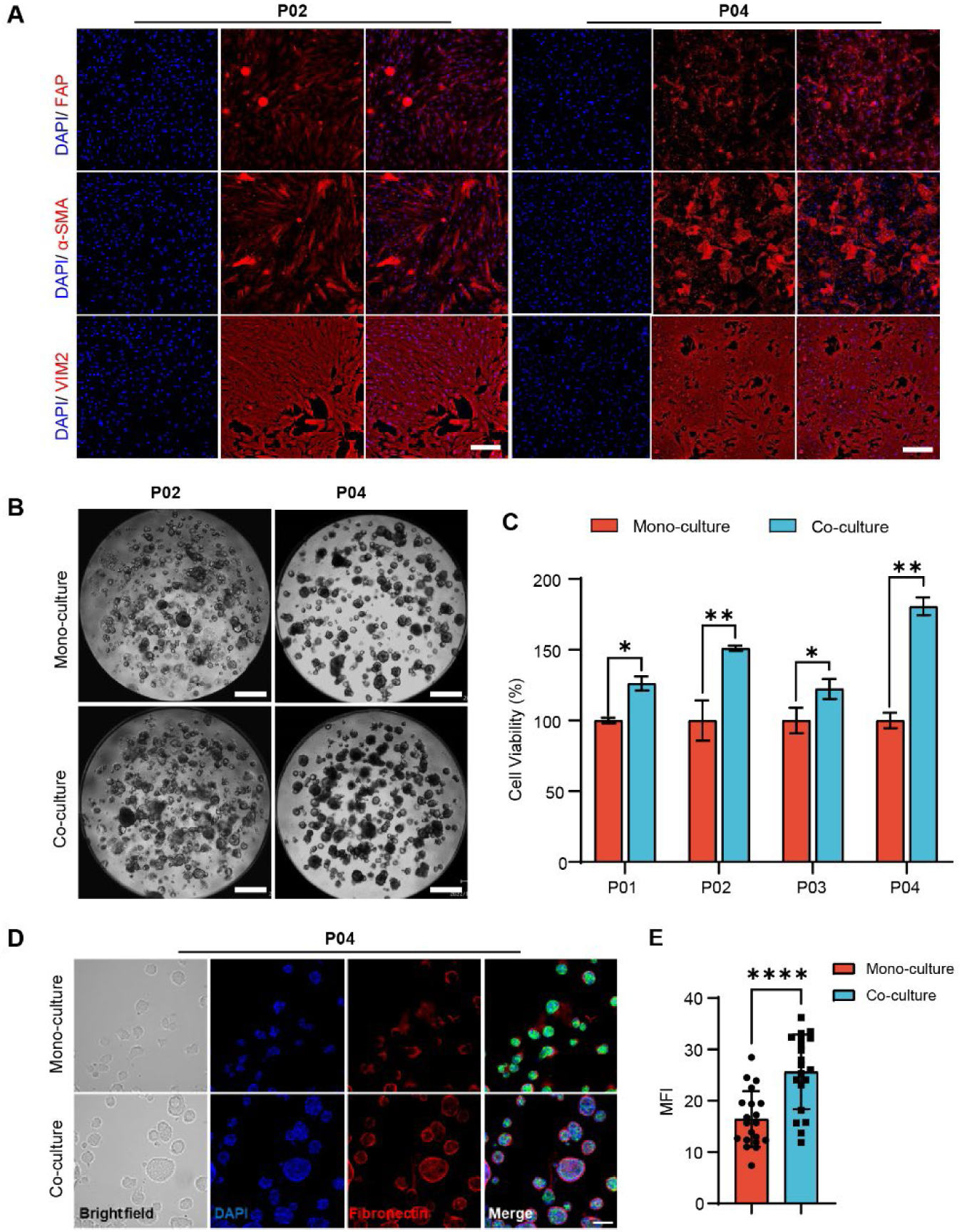
Characterization of CAFs Morphological and Promoting Function by Quantifying PDOs Growth and Epithelia-Mesenchymal Transition Marker. A. The immunofluorescence staining images of FAP, α-SMA, and VIM2 in CAFs from two patients. Scale bars are 100μm as indicated. B. Bright field photos of PDOs in mono-culture and co-culture models. Scale bars are 200μm as indicated. C. The statistical histogram of cell viability in mono-culture and co-culture models. D. Comparative bright field and immunofluorescence staining of Fibronectin (red) and F-actin (green) in mono-culture and co-culture models. Scale bars are 50μm as indicated. E. The statistical mean immunofluorescence intensity of Fibronectin per organoid in Figure 4D. (n≥20) (****P < 0.0001.)

### 2.5 CAFs influence the transcriptome and secretory profile of the co-culture system

To further explore the molecular mechanisms by which non-contact CAFs affect PDO growth and phenotype, we performed transcriptome sequencing and comparative analysis on mono-cultured and co-cultured PDOs. Heatmaps of differentially expressed genes (DEGs) from two donors show significant differences between the two models (Fig. 5A). The DEGs were primarily enriched in pathways such as MAPK, TNF-α signaling, and cytokine-cytokine receptor interaction, suggesting that CAFs affect PDOs mainly through cancer development, inflammation, and microenvironmental interactions (Fig. 5B).

**Figure 5:**
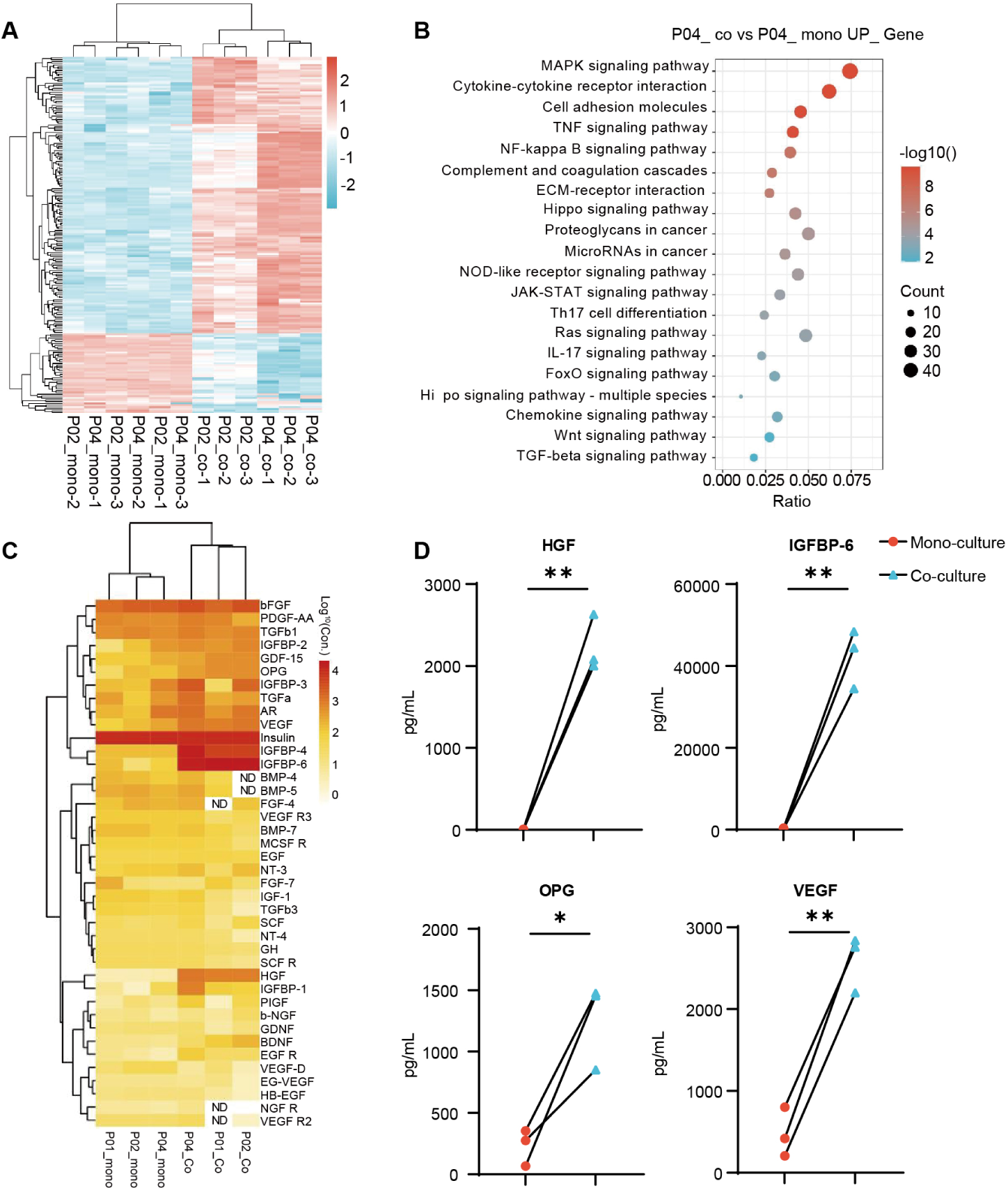
CAFs Influence the Transcriptome and Secretory Profile of Co-culture System. A. Heatmap showing the partial DEGs of PDOs between mono-culture and co-culture models. (n=3) B. Dot plot showing the GO analysis result of up-regulated DEGs in mono-culture and co-culture models of Patient 04. C. Heatmap depicting differentially expressed secreted factors in mono-culture compared with co-culture models. (ND means no detected, n=3) D. The line chart of significantly differentially expressed HGF, IGFBP-6, OPG, and VEGF in co-culture compared with mono-culture models. (n=3) (*P < 0.05, **P < 0.01.)

To elucidate molecular changes in PDOs under co-culture, we measured 40 cytokines in the supernatant from mono-culture and co-culture models of three donors. The heatmap shows that levels of these factors were significantly higher in the co-culture model (Fig. 5C). Eight representative growth factors were significantly increased in the co-culture model across all three donors (Fig. S7A). In addition, HGF, IGFBP-6, OPG, and VEGF showed significant differences (Fig. 5D). These findings are consistent with the reported secretory phenotype of CAFs. Collectively, these results indicate that co-culture markedly reshapes both the transcriptomic landscape and the secretory profile, upregulating key pathways and cytokines that may enhance cellular crosstalk.

### 2.6 Co-culture with CAFs affects drug resistance and recurrence in PDOs

Two representative donors were selected to study the effect of CAFs on tumor drug resistance (Fig. 6). The combination of 5-FU and oxaliplatin was used to assess the drug responses of PDOs and CAFs in co-culture models. For donor P02, the IC50 values of both drugs increased more than 3-fold in the co-culture model compared with mono-culture (5-FU: from 2.66 μM to 9.44 μM; oxaliplatin: from 0.02 μM to 0.06 μM), and the inhibition rates differed significantly at all four high concentrations (Fig. 6A-B). These results suggest that CAF co-culture promotes drug resistance in P02 PDOs. For donor P04, the difference in IC50 between mono-culture and co-culture was within 2-fold (5-FU: 1.95 vs. 3.06 μM; oxaliplatin: 0.22 vs. 0.03 μM), and no significant differences were observed at any concentration, indicating that CAF co-culture did not significantly affect drug sensitivity in P04 PDOs.

**Figure 6:**
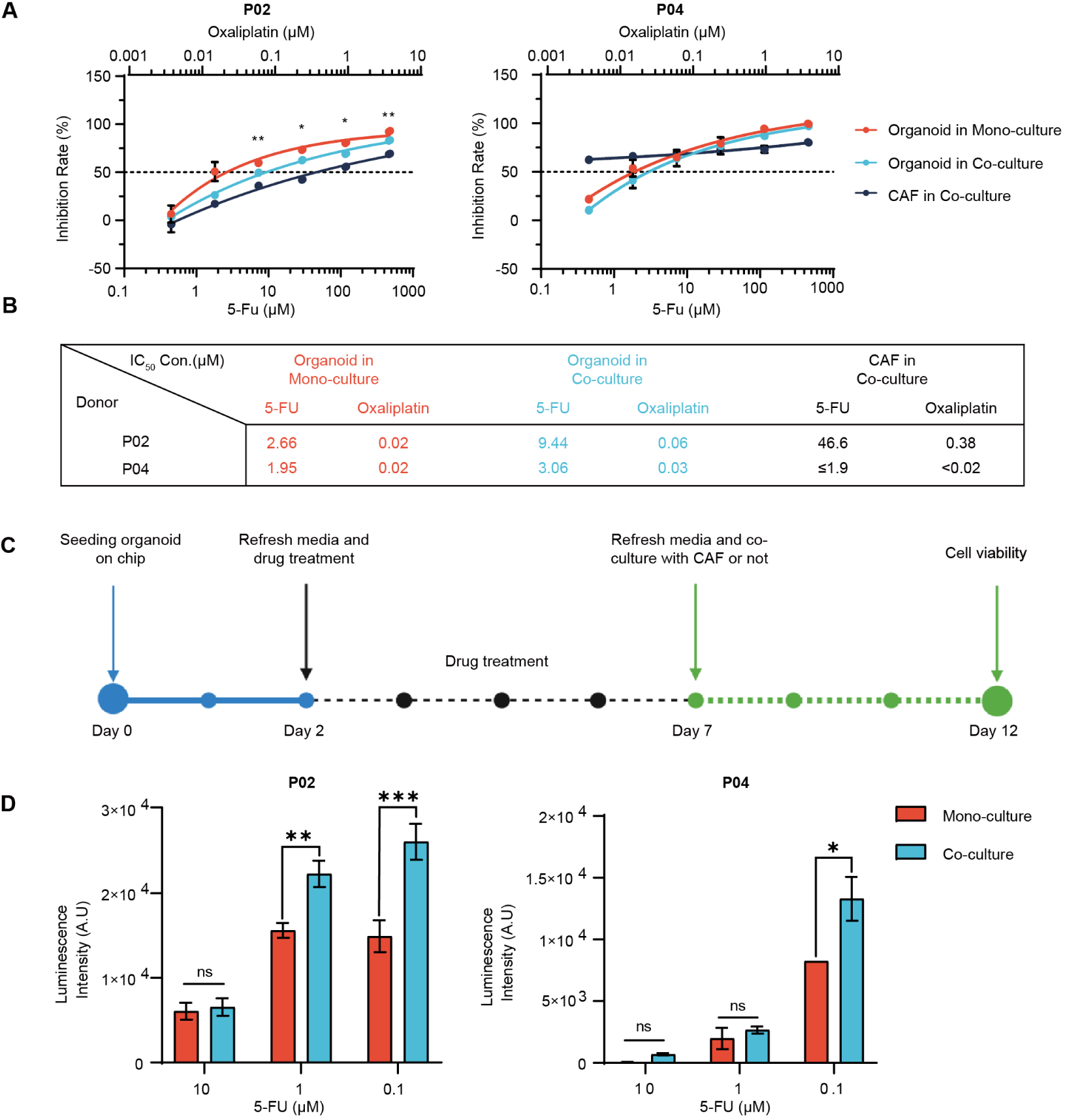
Co-culture with CAFs Affects Drug Sensitivity and Recurrence in PDOs. A. Two patients’ dose-response curves of 5-FU combined with Oxaliplatin in CAF (black), organoid (blue) in co-culture models, organoid (red) in mono-culture models. All data are presented as means ± SD of three replicates. B. The table of different IC50 values in different models with Figure 6A. C. Schematic timeline of the experimental setup to explore the effect of CAFs on organoids recovery after drug treatment. D. The comparison of cell viability of PDOs in mono-culture and co-culture models after drug treatment. (*P < 0.05, **P < 0.01, ***P < 0.001, ns (no-significant) > 0.05)

To determine whether donor-dependent CAF properties influence tumor drug resistance, we examined the sensitivity of CAFs themselves to the chemotherapeutic agents. CAFs from donor P02 had IC50 values of 46.6 μM (5-FU) and 0.38 μM (oxaliplatin), whereas CAFs from donor P04 had IC50 values ≤1.9 μM and <0.02 μM, respectively. Thus, CAFs from P02 were more resistant to the chemotherapy regimen, and this resistance correlated with reduced drug sensitivity of the co-cultured PDOs.

Next, we studied the effect of CAF co-culture on PDO recurrence after drug treatment. PDOs were treated with drugs for 5 days, after which the drugs were withdrawn to mimic tumor recurrence following chemotherapy in vivo (Fig. 6C). Co-culture with CAFs accelerated the recovery and growth of PDOs after treatment with low-dose chemotherapeutic drugs (Fig. 6D and S8). Compared with the mono-culture group, cell viability and density were significantly increased, suggesting that CAFs promote tumor recurrence after drug treatment. These findings indicate that combining CAF-targeted therapies with chemotherapy could counteract tumor drug resistance.

### 2.7 Co-culture Model Accurately Predicts Clinical Efficacy of Chemotherapeutic Agents

Dose-response curves of 24 colorectal cancer (CRC)-derived PDOs treated with the combination of 5-fluorouracil (5-FU) and oxaliplatin were generated (Fig. 7A). Based on the IC50 trichotomy method, PDOs were stratified into three distinct response categories: sensitive (P11, P18), moderately responsive (P14, P21), and resistant (P03, P08), with cutoff values established at 3.01 μM and 7.225 μM, respectively (Fig. 7B). Notably, discordant drug sensitivity profiles were observed between co-culture and mono-culture models for patient P02, with the co-culture model yielding an IC50 of 9.44 μM (resistant) and the mono-culture model producing an IC50 of 2.66 μM (sensitive). In contrast, patient P04 demonstrated concordant results across both models, with IC50 values of 3.06 μM (co-culture) and 1.95 μM (mono-culture), both falling within the moderately sensitive range. To prospectively validate these *in vitro* findings, we monitored the clinical outcomes of patients P02 and P04 following standard chemotherapy. Patient P02 received capecitabine plus oxaliplatin, and computed tomography (CT) imaging at 6 weeks post-treatment revealed progressive disease (PD), confirming clinical drug resistance (Fig. 7C). This clinical outcome was consistent with the co-culture model prediction but inconsistent with the mono-culture model. Patient P04 was treated with capecitabine, oxaliplatin, and bevacizumab, and CT evaluation at 16 weeks demonstrated partial response (PR) (Fig. 7D), with the clinical response aligning with predictions from both co-culture and mono-culture models.

**Figure 7:**
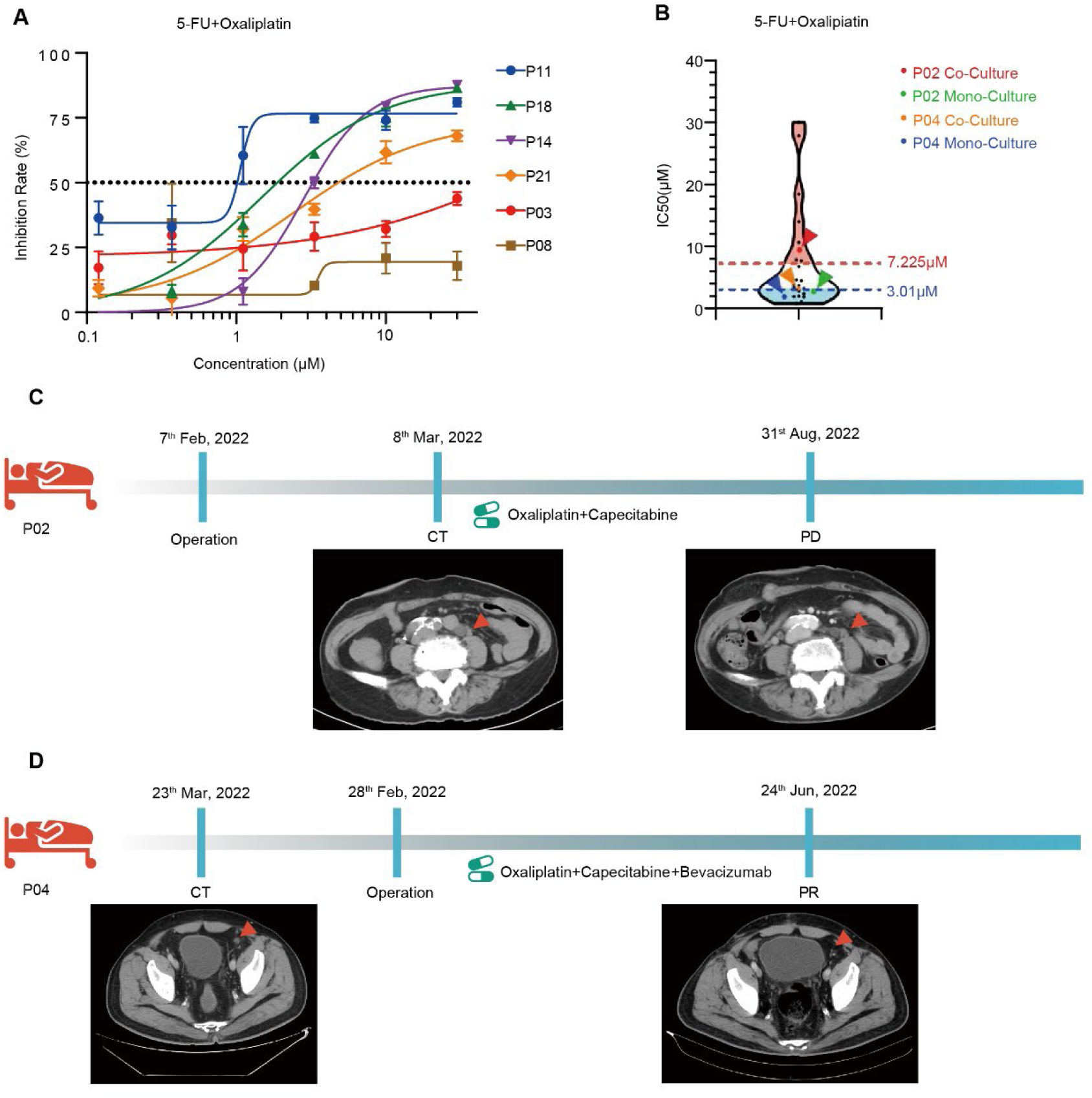
Drug Sensitivity Testing to Combined 5-FU and Oxaliplatin and Clinical Concordance Analysis. A. Respective dose-response curves of 5-FU combined with Oxaliplatin in organoids models from 6 donors. B. Violin plot showing the distribution of IC50 values of 5-FU in the 24 organoid cells, and IC50 values received by patients P02 and P04. C. Clinical and radiographic follow-up data of patients P02. D. Clinical and radiographic follow-up data of patients P04.

To further assess the predictive accuracy of the co-culture model for alternative chemotherapeutic agents, drug sensitivity assays were performed using irinotecan (CPT-11) and raltitrexed across 18 PDOs, with response cutoff values defined at 1.63 μM and 6.47 μM using the IC50 trichotomy method (Fig. S9B). Under this classification, the P07 PDO co-culture model exhibited differential sensitivity to the two regimens: the 5-FU/oxaliplatin combination fell within the resistant range (IC50 = 14.42 μM), whereas the CPT-11/raltitrexed regimen was classified as sensitive (IC50 = 0.89 μM) (Fig. S9A and S9B). Clinically, patient P07 presented with liver metastases from CRC and was initially treated with oxaliplatin plus capecitabine; CT imaging at 10 weeks showed disease progression, indicating resistance to this regimen. Following a switch to irinotecan, raltitrexed, and bevacizumab, subsequent CT evaluation revealed stable disease in the liver metastases. Collectively, these findings demonstrate that co-culture models provide superior predictive accuracy for clinical treatment sensitivity compared to mono-culture models.

## 3. Discussion

This study aimed to explore the role of CAFs in the tumor microenvironment and their impact on tumor growth, tumor drug resistance and recurrence by using a dynamic co-culture chip system. Here, we discuss the implications of our findings and how they advance the understanding of tumor-stromal interactions and personalized medicine. Different cell types were seeded in separate regions of the chip for non-contact co-culture, enabling cell communication via the paracrine pathway, and the nested design allowed convenient medium exchange without disruption of the 3D organoids. The open-top design also allowed easy access for changing the culture medium and for drug administration. Compared with traditional co-culture platforms such as 24-well plate or Transwell, BAC OC/OD chip is more suitable for 3D cell culture and detection, and demonstrate strong superiority in terms of accommodating cell types, mass transfer efficiency, convenient liquid exchange operation, real-time imaging and cell recovery. Our newly developed dynamic co-culture chips offer a biomimetic platform that enables the study of CAFs and tumor organoids under conditions that closely resemble the in vivo tumor microenvironment. The comparison between static and dynamic culture conditions demonstrated the superiority of the latter in promoting the growth of organoids and cell-cell interaction. The design allows for non-contact co-culture, which is crucial for studying the paracrine effects of CAFs on tumor organoids highlighting the importance of mimicking in vivo conditions for accurate preclinical studies.

Our results show that CAFs significantly promote the growth, inflammation and upregulation of EMT marker of PDOs in a non-contact co-culture model. The results suggest that a variety of soluble factors secreted in the co-culture system play a crucial role in tumor progression and indicate that CAFs may induce EMT and cell signaling. These findings underscore the influence of the tumor microenvironment on therapeutic outcomes. Notably, the co-culture model was able to predict the clinical response to chemotherapy more accurately in patients, suggesting that it could serve as a valuable tool for personalized medicine. Moving forward, larger clinical studies are warranted to further validate the predictive power of the co-culture model. The finding that CAFs promote tumor recurrence after drug treatment highlights the need to consider the microenvironment in developing therapeutic strategies aimed at preventing relapse. Additionally, exploring the combination of targeted therapies against CAFs with conventional chemotherapy holds promise for overcoming drug resistance and improving treatment outcomes in colorectal cancer.

In conclusion, our study provides a novel tool for establishing co-culture models to inform personalized treatment strategies. And provides compelling evidence for the significant role of CAFs in modulating the behavior of colorectal cancer cells. Further research is needed to fully elucidate the complex interactions within the tumor microenvironment and to translate these insights into more effective cancer therapies.

## 4. Experimental Section

### Ethics statement

This study was approved by the Ethics Committee of the National Cancer Center/Cancer Hospital, Chinese Academy of Medical Sciences (NCC2022C-431). Written informed consent was obtained from all patients providing colorectal cancer samples. All samples were de-identified before use. The research was performed in accordance with the Declaration of Helsinki.

### Establishment and culture of CRC tumor organoids

Fresh gastrointestinal tumor tissue was washed 5–10 times with PBS containing penicillin-streptomycin, then cut into fragments no larger than 2 mm³. The tissue was digested with Tissue Enzyme Solution Ⅱ (KS100130, Daxiang Biotech) using Intelligent Tissue Dissociator (D8, Daxiang Biotech) under 37°C for 10-15 min, followed by mechanical dissociation through repetitive pipetting in DMEM/F12. The supernatant from enzymatic digestion was filtered through a 70 µm cell strainer, centrifuged, and the pellet was resuspended in Matrigel (MG100101, Daxiang) and seeded into 24-well plates for culture. After polymerization at 37 °C for 10 minutes, the organoid dome was overlaid with 500 µL of organoid medium (OC105131, Daxiang), which was renewed every 2-3 days. Organoids were passaged at a ratio of 1:2 to 1:5 depending on growth. During passaging, organoids were collected with DMEM/F12, centrifuged, digested with Organoid Digestion Solution (100145, Daxiang Biotech) for 3–5 minutes to dissociate them, and then re-seeded into Matrigel.

### Isolation and culture of autologous CAFs

After enzymatic digestion of the tissue with collagenase, the resulting precipitate was collected, and the tissue fragments were evenly distributed onto the bottom of a T25 cell culture flask, then incubated for 15–30 minutes. Subsequently, 2.5 mL of CAF medium (FC100101, Daxiang) was slowly added to the flask, and the medium was changed every three days. When CAF confluence reached approximately 80%, cells were passaged. Cells were passaged at least once before freezing and used for experiments between passages 2 and 10. CAF medium consisted of 1640 medium supplemented with 1% penicillin-streptomycin and 10% FBS.

### Establishment of mono-culture and CAF co-culture organoid models

To establish organoid mono-culture and CAF co-culture models on the BAC-OC and BAC-OD chips, 6 mL of sterile water was first added to the volatile protection chamber of the chip, which was then placed in an incubator to equilibrate. Before use, chips were removed from the incubator and allowed to reach room temperature. Organoids typically grew for 5–7 days to reach passaging density, while CAFs were grown to approximately 80% confluence before digestion and seeding.

For the organoid mono-culture model, organoids were digested and prepared as a single-cell suspension containing a small proportion of clusters. The cell concentration was adjusted to 500 cells/µL with organoid culture medium and kept on ice. An organoid-Matrigel suspension at 200 cells/µL was prepared by mixing the organoid suspension with Matrigel at a 2:3 ratio. Next, 50 µL and 5 µL of this suspension (containing 10,000 and 1,000 organoids, respectively) were seeded into the organoid loading chambers of the BAC-OC and BAC-OD chips. The chips were then incubated at 37 °C for 10 minutes to allow solidification, after which organoid medium containing 0.1% anti-apoptotic agents was added.

For the co-culture model, CAFs were harvested by digestion, and the cell density was adjusted with complete CAF culture medium. A CAF-Matrigel suspension at 1000 cells/µL was prepared by mixing the CAF suspension with Matrigel. Then, 100 µL and 20 µL of this Matrigel mixture (containing 100,000 and 20,000 CAFs, respectively) were seeded into the CAF loading chambers and incubated at 37 °C for 15 minutes to solidify. Each well was then carefully supplemented with CAF culture medium to avoid disturbing the cell-Matrigel suspension. After 2–3 days of mono-culture for organoids and CAFs, co-culture was initiated by connecting the two chambers. A co-culture medium containing 50% CAF medium and 50% organoid medium (250 µL for BAC-OD and 1000 µL for BAC-OC per co-culture unit) was added directly to each co-culture unit.

On the BAC-OC chip, 50 µL of hydrogel-cell mixture was seeded into the cell culture part of the organoid culture well, and 100 µL of hydrogel-cell mixture into the CAF culture well. For medium storage, 200 µL of culture medium was added to both the organoid and CAF culture wells. For a three-co-culture unit, a total of 3 mL of culture medium was added to connect the organoid and CAF culture wells.

On the BAC-OD chip, 10 µL of hydrogel-cell mixture was seeded into the organoid culture well and 20 µL into the CAF culture well. For medium storage, 50 µL of culture medium was added to each well. To connect a single co-culture unit, 250 µL of culture medium was added to connect the organoid and CAF wells. For a three-co-culture unit, 1.5 mL of medium was added to connect three organoid wells and their corresponding CAF wells.

*Evaluation of drug efficacy on the BAC-OD chip:* After 1-2 days of co-culture, drug efficacy testing was performed. First, the culture medium was aspirated from the chip platform. The medium in the co-culture wells was carefully removed using a pipette, and 370 µL of drug solution was added for incubation. The cells were cultured for an additional 3-5 days before testing. The drug regimen consisted of 5-FU and oxaliplatin (1:1 ratio). The maximum concentrations were 462 µM for 5-FU and 3.8 µM for oxaliplatin, followed by 4-fold serial dilutions to generate a total of 6 concentration gradients. Co-cultures were treated with drugs for 5 days, and a CTG assay was performed on the final day to assess the effects.

### Organoid cell recovery after drug treatment

On day 0, organoids were seeded into the organoid loading chamber of the microchip and allowed to stabilize for two days. On day 2, organoids were treated with 5-FU and oxaliplatin (1:1) at a maximum concentration of 462 µM for 5-FU and 3.8 µM for oxaliplatin, followed by 10-fold serial dilutions to generate three concentration gradients. On day 5, CAFs were seeded into the CAF loading chamber. After two days of stabilization, CAFs and organoids were co-cultured starting on day 7, and the co-culture medium was changed to a 1:1 mixture of CAF medium and organoid medium. Continuous observation was performed for 5 days. CTG assays were performed on days 10–12 to assess the effects of drug treatment on organoids.

### Viability assessment of organoids

Organoid viability was measured using the CellTiter-Glo 3D Cell Viability Assay (Promega) according to the manufacturer’s instructions. Briefly, the CellTiter-Glo reagent and organoid culture medium were mixed at a 1:1 volume ratio. Luminescence was detected using a multiplate reader. Dose-response curves and IC50 values were determined based on the cellular response to the drug. The inhibition rate was calculated using the following equation: Inhibition rate (%) = (1 − fluorescence with drug administration / fluorescence without drug administration) × 100.

### Immunofluorescence staining

Organoids were characterized by immunofluorescence staining for CK20 (1:200, 13063S, CST), CDX2 (1:100, 12306S, CST), Ki67 (1:1000, 9449, CST), and F-actin (1:100, CA1640, Solarbio). CAFs were characterized by staining for Vimentin (1:200, 9856S, CST), FAP (1:100, 66562S, CST), and α-SMA (1:50, 34105S, CST). Organoids and CAFs were cultured for 5–7 days and then fixed with glutaraldehyde for 30 minutes at room temperature. Fixed cells were permeabilized with 0.5% Triton X-100 in PBS for 30 minutes at room temperature and blocked with 5% bovine serum albumin (BSA) for 60 minutes after washing. Organoids and CAFs were incubated overnight at 4 °C with unconjugated primary antibodies, then incubated with secondary antibodies conjugated with F488 (1:400, A11008, Invitrogen) or AF594 (1:1000, 8889S, CST) for 1.5 hours in the dark. Directly labeled antibodies were incubated at room temperature for two hours. After incubation, cells were washed three times with PBS for 5 minutes each and stained with DAPI (1:200, C0060, Solarbio) for nuclear visualization. Final imaging analysis was performed using an ImageXpress Confocal HT.ai (Molecular Devices, San Jose, CA, USA).

### Hematoxylin and eosin (H&E) and immunohistochemistry staining

PDOs were first recovered from Matrigel using Cell Recovery Solution (Corning). PDOs and parental tissues were fixed with 4% freshly prepared paraformaldehyde at 4 °C for 24 hours, dehydrated, embedded in paraffin blocks, and sectioned into 5-µm slices. These sections were subjected to standard H&E and immunohistochemical staining. For H&E staining, sections were stained with hematoxylin and eosin. For immunohistochemistry, sections were stained with primary antibodies against Ki-67, CDX2, and CK20 as described above. H&E and immunohistochemical images were acquired using an inverted microscope (Olympus).

### RayBiotech cytokine microarray analysis

The Quantibody® Human Growth Factor Array 1 (QAH-GF-1, RayBiotech Inc.) was used to measure the supernatant levels of 40 growth factors in two groups, each containing three samples. The experimental procedure followed the manufacturer’s instructions. Briefly, glass slides were equilibrated to room temperature and air-dried for 1–2 hours. The lyophilized cytokine standard mix was reconstituted, and a series of dilutions was prepared. Slides were blocked with sample diluent for 30 minutes at room temperature. Standards or samples were added to each well and incubated overnight at 4 °C. After washing with Wash Buffers I and II, a biotinylated antibody cocktail was added to each well and incubated for 1–2 hours at room temperature, followed by another wash. Streptavidin conjugated to a Cy3-equivalent dye was added and incubated in the dark for 1 hour at room temperature, followed by a final wash with Wash Buffer I. Slides were then dried and imaged using a compatible laser scanner, and data were analyzed with Q-Analyzer software.

### Whole-exome sequencing (WES) analysis

PDOs were harvested, and DNA was extracted. Samples were sent to Genetron Health (Beijing) Co., Ltd. for whole-exome sequencing analysis using an Illumina HiSeq platform. Sequence reads were aligned to the human reference genome GRCh37 using BWA-MEM (v0.7.10), which employs the Burrows-Wheeler Aligner with maximal exact matches. BAM files were processed to remove duplicate reads. Somatic variants were identified by providing reference and tumor/organoid sequencing data to MuTect (v3.1-0-g72492bb) with default parameters. Effect predictions and annotations were added using ANNOVAR. To detect somatic copy number alterations (CNAs), BAM files were analyzed for read-depth variations using Control-FREEC (V9.1) by comparing tumors or organoids to adjacent normal tissue. Mutational signatures were analyzed using the R package BSgenome.

### RNA-seq data analysis

The reference genome and corresponding gene model annotation files were obtained directly from the genome website. An index of the reference genome was constructed using Hisat2 v2.0.5, and paired-end clean reads were aligned to the reference genome using the same software. The mapping results were then analyzed with featureCounts v1.5.0-p3 to quantify transcript abundance. Fragments per kilobase of transcript per million mapped reads (FPKM) were calculated for each gene based on both gene length and the number of reads mapped to that gene. Differential expression analysis was performed using DESeq2, which provides statistical routines for identifying differentially expressed genes based on a negative binomial distribution model. Resulting P-values were adjusted using the Benjamini-Hochberg method to control the false discovery rate.

### Functional enrichment analysis

A curated list of differentially expressed genes (fold change > 2) was submitted for analysis. This gene set was compared against a comprehensive background gene list for Homo sapiens. Using the Gene Ontology (GO) framework, the Biological Process category was specifically selected to annotate and elucidate the biological functions significantly associated with the differential gene expression. Enrichment scores, P-values, and false discovery rate (FDR) adjustments were calculated to evaluate the significance of the identified functional categories. Pathways with an FDR-adjusted P-value < 0.05 were considered statistically significant, and the top enriched terms were visualized using dot plots or bar graphs for clearer interpretation of the data.

### Statistical analysis

To determine whether differences between two experimental groups were significant, a t-test was used. A P-value threshold of 0.05 was established; P > 0.05 was interpreted as indicating no significant difference, whereas P < 0.05 was considered statistically significant. All data analyses were conducted using GraphPad Prism version 8, which facilitated calculation of means ± standard error for each group, as well as generation of histograms and box plots for visual representation of data distribution.

## Acknowledgements

1. R. Xiao, Z. Wang and Y. Zhou contributed equally to this work. This study was financially supported by National Key Research and Development Project (2023YFC3505000), Beijing Municipal Science & Technology Commission (Z231100007223001), National Natural Science Foundation of China (82422076 and 82174086), Scientific and Technological Innovation Project of China Academy of Chinese Medical Sciences (C12024C002YN), and Key Research and Development Program of Ningxia (2024BEG01006).

## Data Availability Statement

The data that support the findings of this study are available from the corresponding author upon reasonable request.

---

**Supplementary Figure 1:**
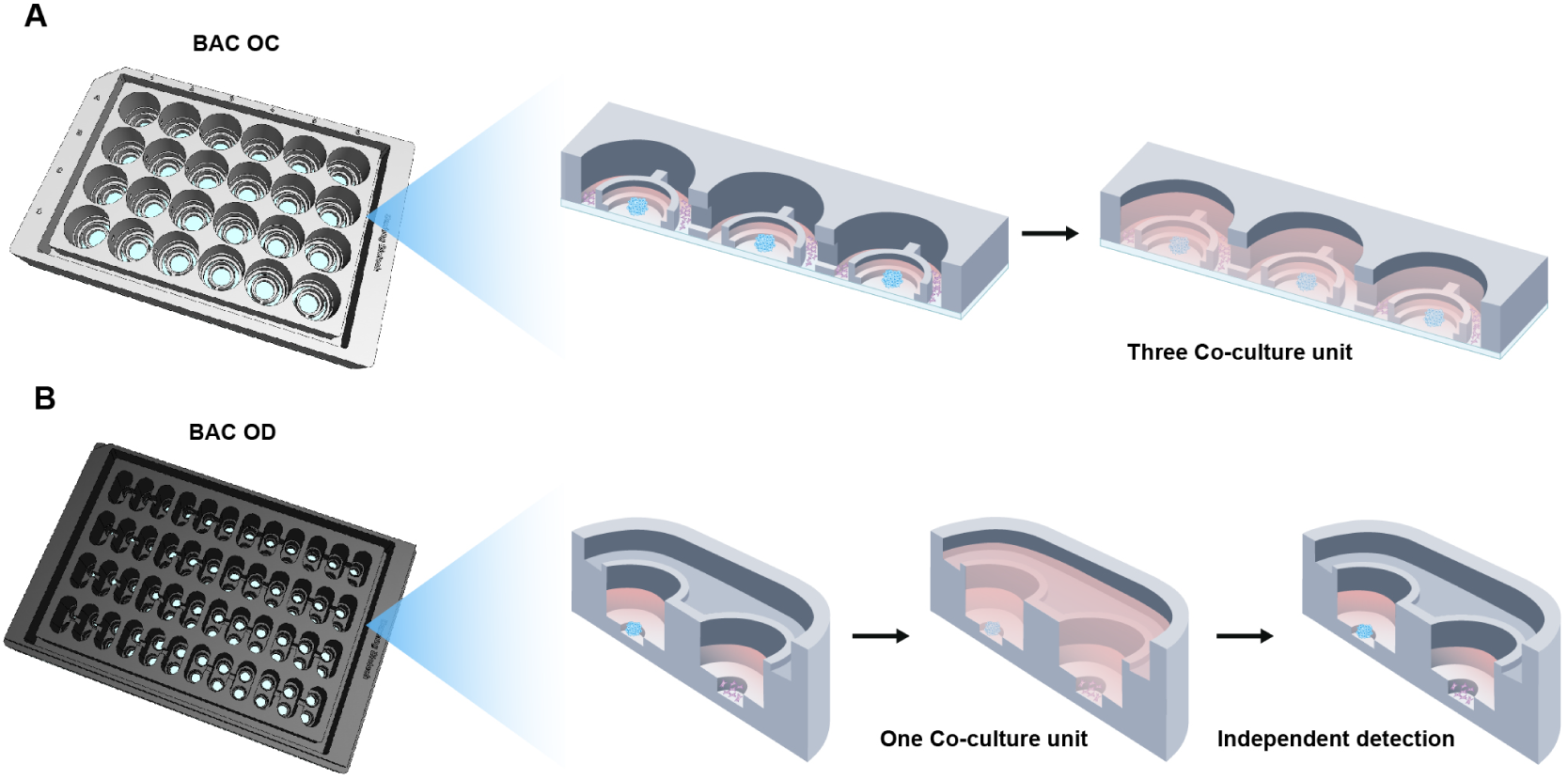
Introduction to the Co-culture Usage modes of the Chips. A. The Co-culture usage modes of the BAC OC Chip. B. The Co-culture usage modes of the BAC OD Chip.

**Supplementary Figure 2:**
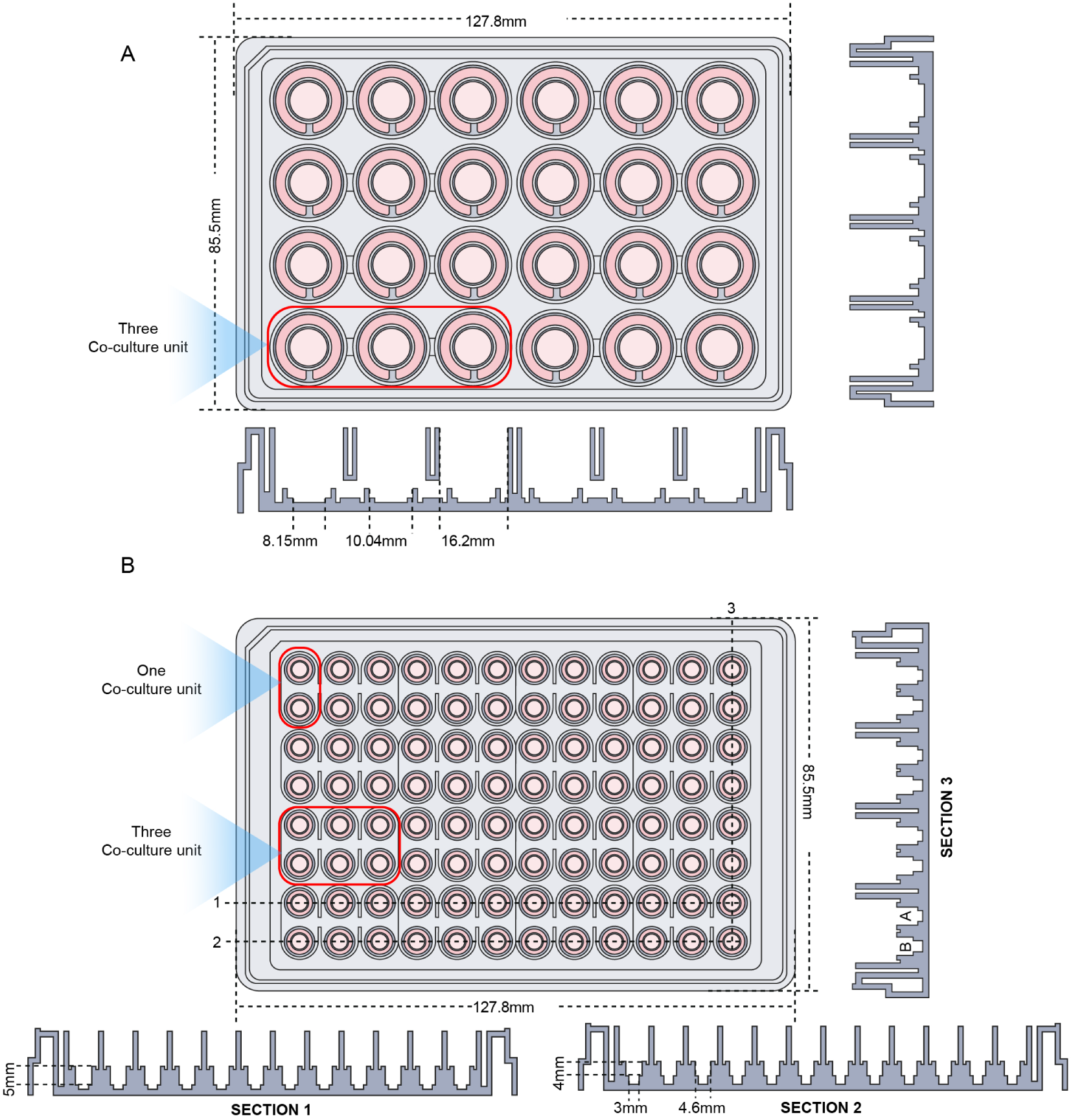
Design Details of Dynamic Co-culture Chips BAC OC and BAC OD. A. Design Details of Dynamic Co-culture Chips BAC OC. B. Design Details of Dynamic Co-culture Chips BAC OD.

**Supplementary Figure 3:**
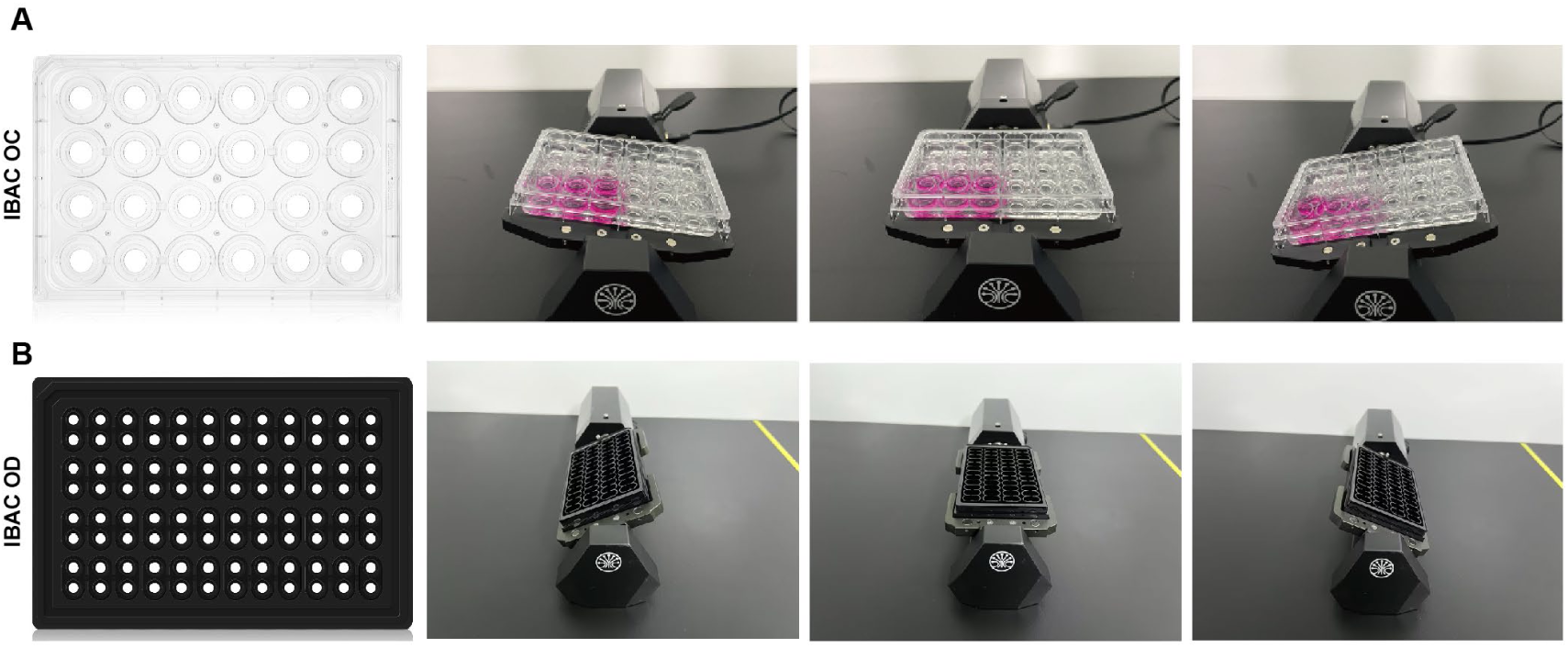
The Images of Dynamic Culture Using BAC OC and BAC OD.

**Supplementary Figure 4:**
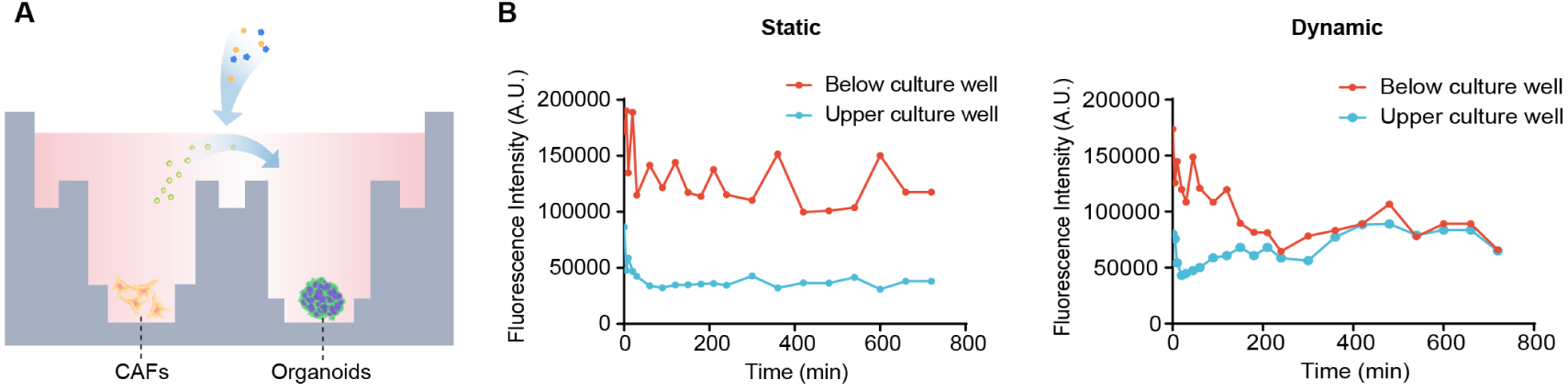
Characterizing the Molecular Diffusion Efficiency of Dynamic and Static Models in BAC OD. A. The schematic diagram of the molecular transfer process in adjacent wells in a module. B. Quantifying the transfer efficiency of FITC-dextran in adjacent wells in dynamic and static culture models.

**Supplementary Figure 5:**
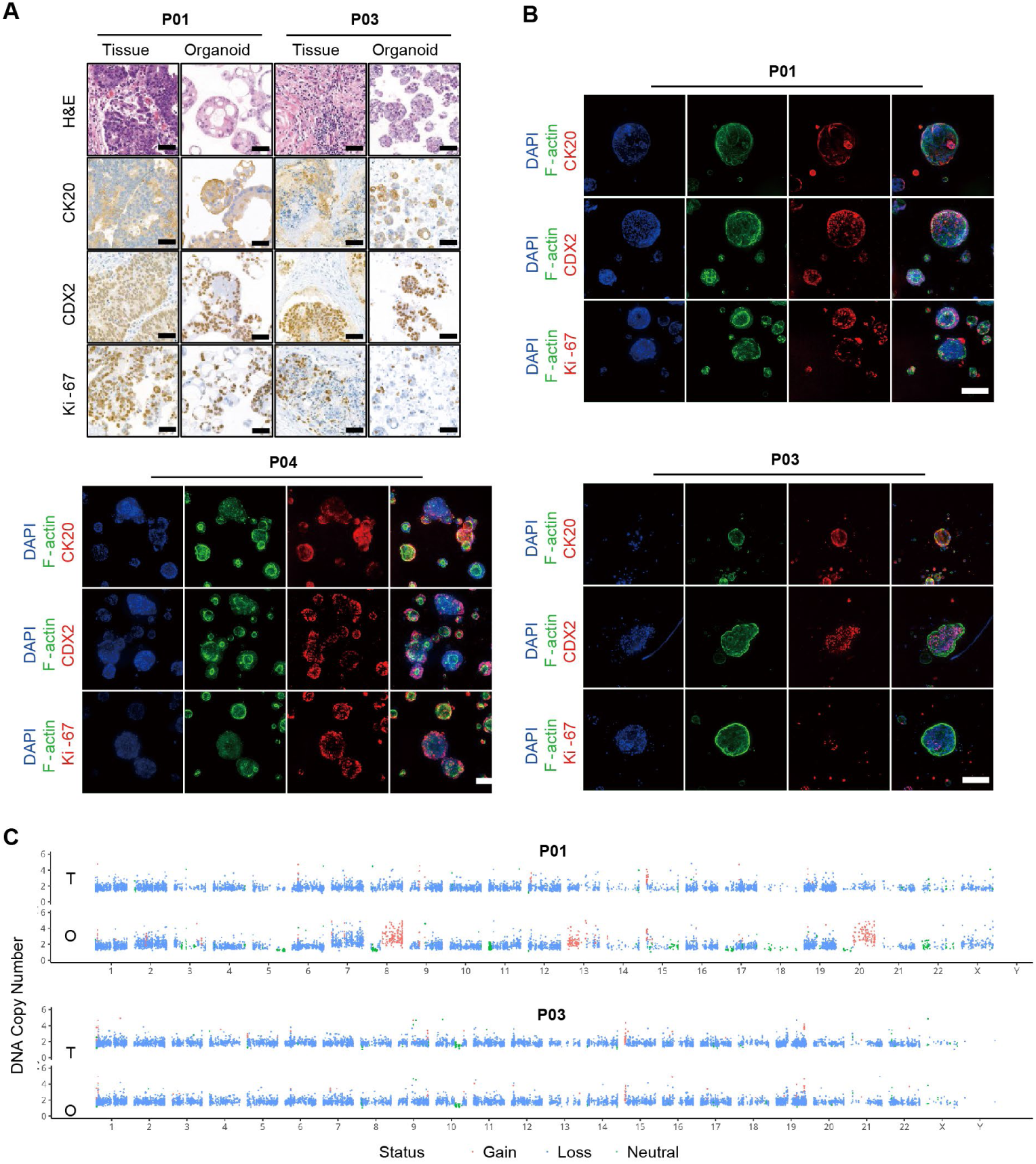
PDOs are Highly Consistent with Patient Tissues. A. Comparative H&E staining and immunohistochemical staining images of PDOs and parental tumor tissues. The typical markers are CK20, CDX2, and Ki-67. Scale bars are 50μm as indicated. B. The images of immunofluorescence staining of PDOs. The typical markers are CK20, CDX2, and Ki-67. Scale bars are 100μm as indicated. C. Scatterplots illustrating genome-wide gene CNVs of the PDOs and parental tumor tissues (red: gain; blue: loss; green: neutral).

**Supplementary Figure 6:**
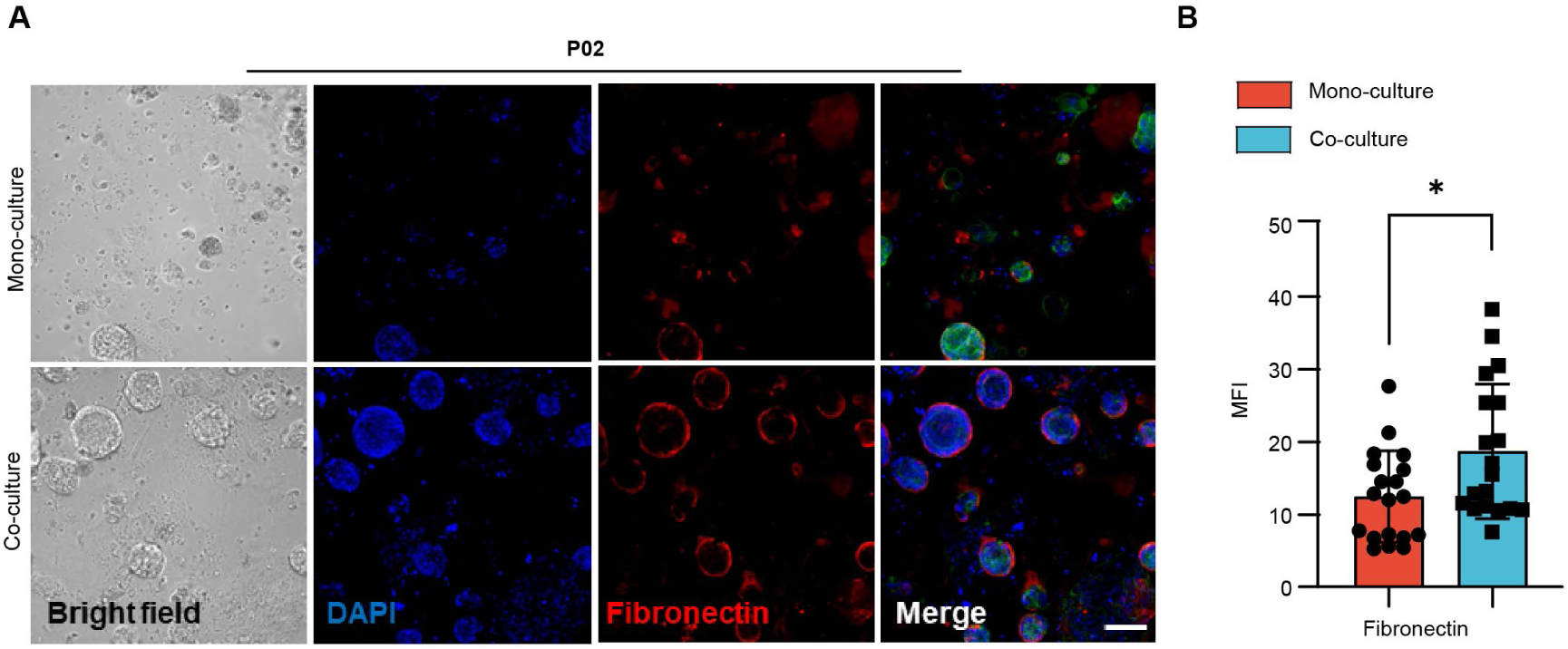
Co-culture with CAFs Promotes PDOs Epithelial-Mesenchymal Transition. A. Comparative bright field and immunofluorescence staining of Fibronectin (red) and F-actin (green) in mono-culture and co-culture models. Scale bars are 50μm as indicated. B. The statistical mean immunofluorescence intensity of Fibronectin per organoid in Supplementary Figure 6A. (n≥20) (*P < 0.05)

**Supplementary Figure 7:**
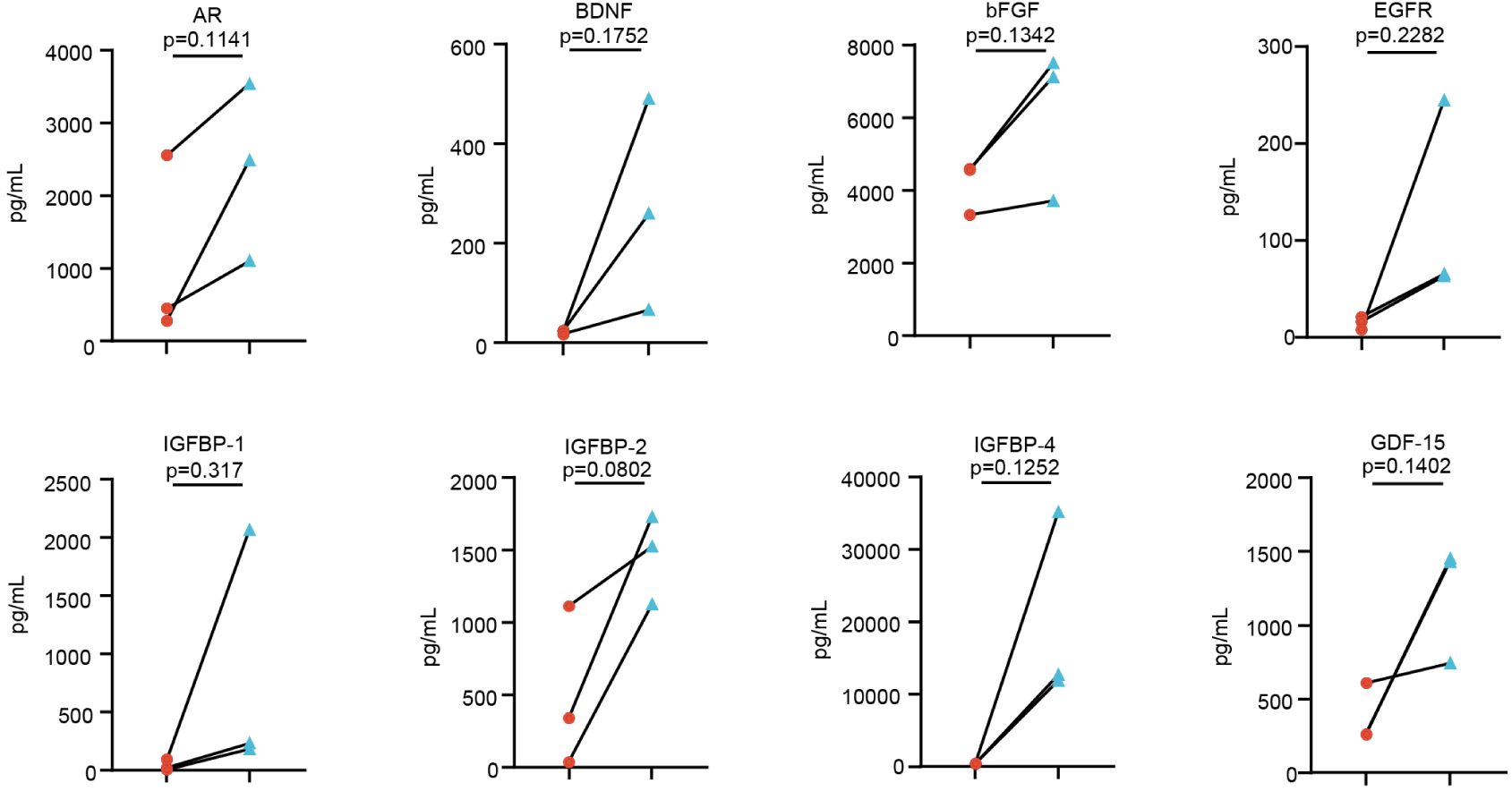
Secretory Factors with Up-regulated Expression in Co-culture Model. Line charts of up-regulated expression of HGF, IGFBP-6, OPG, and VEGF in co-culture compared with mono-culture models.

**Supplementary Figure 8:**
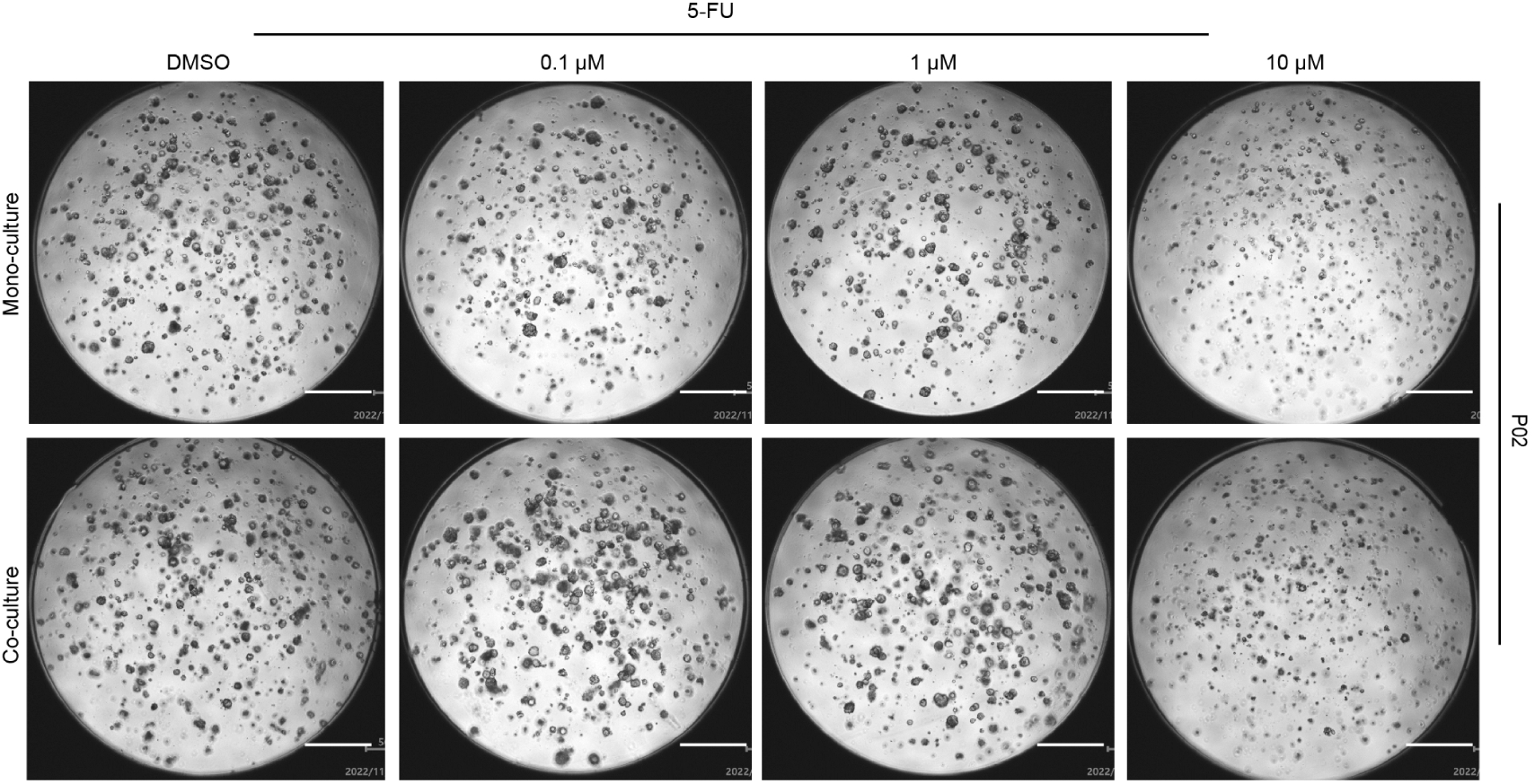
Response to Different Drug Concentrations in Mono-culture and Co-culture Models. Bright field photos of PDOs’ response to different drug concentrations in mono-culture and co-culture models.

**Supplementary Figure 9:**
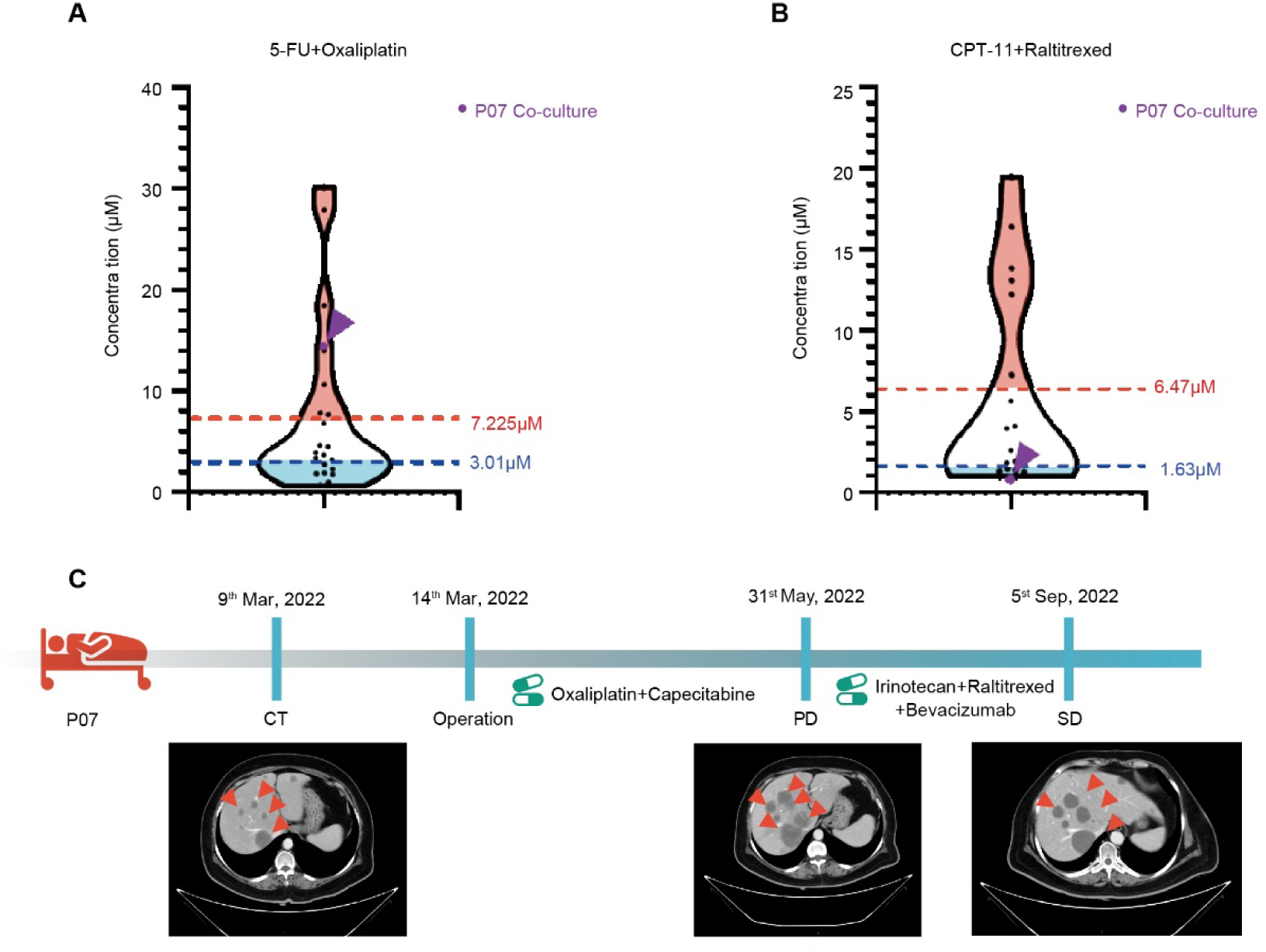
Drug Sensitivity Testing in Models and Clinical Concordance Analysis. A. Violin plot showing the distribution of IC50 values of the combined 5-FU and oxaliplatin in the 24 organoid cells, and IC50 values received by patients P07. B. Violin plot showing the distribution of IC50 values of the combined CPT-11 and raltitrexed in the 18 organoid cells, and IC50 values received by patients P07. C. Clinical and radiographic follow-up data of patient P07.

